# Phase diagrams of bone remodeling using a 3D stochastic cellular automaton

**DOI:** 10.1101/2023.12.21.572505

**Authors:** Anna-Dorothea Heller, Angelo Valleriani, Amaia Cipitria

**Affiliations:** Department of Biomaterials, Max Planck Institute of Colloids and Interfaces, Potsdam, Germany; Group of Bioengineering in Regeneration and Cancer, Biogipuzkoa Health Research Institute, San Sebastian, Spain; IKERBASQUE, Basque Foundation for Science, Bilbao, Spain

**Author notes:** These authors contributed equally to this work (Angelo Valleriani, Amaia Cipitria).

## Abstract

We propose a 3D stochastic cellular automaton model, governed by evolutionary game theory, to simulate bone remodeling dynamics. The model includes four voxel states: *Formation, Quiescence, Resorption*, and *Environment*. We simulate the *Resorption* and *Formation* processes on separate time scales to explore the parameter space and derive a phase diagram that illustrates the sensitivity of these processes to parameter changes. Combining these results, we simulate a full bone remodeling cycle. Furthermore, we show the importance of modeling small neighborhoods for studying local bone microenvironment controls. This model can guide experimental design and, in combination with other models, it could assist to further explore external impacts on bone remodeling. Consequently, this model contributes to an improved understanding of complex dynamics in bone remodeling dynamics and exploring alterations due to disease or drug treatment.

**Author summary:** Functional well-balanced bone remodeling is crucial for a healthy skeleton. Understanding the various causes for the disruption of this complex process requires models that combine behavior on micro- and macro-level. We propose a 3D stochastic cellular automaton model, where each voxel can take one of four different states representing phases of bone remodeling: *Formation, Quiescence, Resorption* and *Environment*. With a qualitative analysis, we show how and when small changes in key interaction parameters can make the difference between healthy and pathological bone remodeling. We also highlight the importance of spatial modeling and the size of the neighborhood for investigating local mechanisms. Our proposed model opens the window to explore the complex interplay of factors essential for healthy bone remodeling and potential alterations due to disease or drug treatment. In the long term this could help with the development of new treatment options for bone diseases.

## Introduction

Bone remodeling is a very complex process necessary to ensure a healthy bone structure. This process can be disturbed by a number of causes, for example hormonal imbalance, extreme load conditions or cancer-related bone disease. To understand this complex interplay, bone remodeling has been extensively studied over the last decades including *in silico* models on macro- and micro-level. Early studies were focused on adaptation of the bone structure and density based on the prevailing loading conditions. Whereas first models were still ran in 2D [1], [2], [3], [4], [5], [6] and [7], later models would move to a 3D domain [8], [9], [10], [11], [12], [13] and [14] partially include experimental data from micro-computed tomography scans [15], [16], [17], [18] and [19].

These models have greatly advanced our understanding of bone architecture. However, they all focus on the (dis-)appearance of mineralized bone as a response to a mechanical stimulus. Moreover, they contain a very simple representation of the many cellular processes underlying the coupling of bone resorption and formation. Whereas the previous models have been very successful in predicting the change of bone macro structure, there was (and still is) a need for micro-level models of multi-cellular interaction. Early attempts of non-spatial models were already made in the 2000s [20], [21] and continued during the following decades [22], [23], [24], [25]. At the same time spatial multi-cellular models were introduced [26], [27], [28], [29], allowing to illustrate the importance of local signaling mechanisms compared to global ones. More recently, some of these models have been coupled with mechanical simulations to investigate the interplay of bone resorption and formation [30], [31], [32].

The results of those models are promising, but most of them (except for [20] and [29]) are limited to specific signaling pathways. Indeed, there are still open questions and ongoing discussions about the exact interplay and origin of signaling cytokines [33], [34], [35], [36]. This discussion becomes especially relevant when bone remodeling is coupled with diseases such as cancer [37]. From the perspective of *in silico* modeling, there is a delicate balance between adding another biochemical or mechanical pathway to better approximate the reality, and creating a model that is too complex to analyze and run, especially when complex differential equations are involved. We believe that a stochastic cellular automaton model that operates on the principles of evolutionary game theory (EGT), as introduced by Ryser et al [29], can help solving this contradiction.

In game theory, in its simplest formulation, each player follows one out of a set of different competing strategies. The outcome of playing one chosen strategy depends on which strategy the opponent is playing and is integrated into a matrix called payoff matrix. In EGT the payoff matrix determines the fitness of an individual based on its strategy and the relative frequency of all other strategies present in the population at a given time. The fitness eventually determines if that individual propagates and, consequently, if the relative frequency of that strategy increases or not [38]. A general question in the context of EGT relates to which conditions allow different strategies to coexist. It has been found that coexistence depends on the neighboring structure of the game: spatially explicit models, in which individuals interact only with their neighbors deliver different and broader coexisting conditions than mean-field models [29], [38].

Cellular automaton models are ideal to recapitulate local dynamics of the microenvironment since they offer a straightforward definition of neighborhood. In this work, we implement a 3D stochastic cellular automaton, where voxels interact only with their nearest neighbors in a volume of interest representing bone tissue. At each time point, each voxel can take one of four different states representing the different phases of bone remodeling: *Resorption, Formation, Quiescence* and *Environment*. To create a compact representation of the frequency-dependent interaction between the local phases of bone remodeling, we consider a voxel as an individual and its state as a strategy in an evolutionary game. This representation encodes knowledge about the mutual impact the main actors of bone remodeling (osteoclasts, osteoblasts, osteocytes and environmental impacts) have on each other. Each payoff parameter in the model therefore is indirectly connected to the biological processes.

We show that a stochastic cellular automaton model governed by EGT rules is capable to simulate interactions of the microenvironment of bone remodeling by using payoff parameters instead of several differential equations. Screening single specific payoff parameters, we investigate parts of the parameter space and highlight phase transitions of the model dynamic. Associated with that, we illustrate why modeling a small neighborhood is essential for investigating local mechanisms. Assigning different payoff parameters to model physiological and pathological behavior of bone, we can draw first conclusions on the necessary interactions of *Resorption, Formation, Quiescence* and *Environment* in healthy bone. By doing so, we can identify key parameter combinations required for physiological bone remodeling, as well as the sensitivity of the system transitioning to pathological remodeling. This model opens the window to explore the complex interplay of factors essential for healthy bone remodeling and potential alterations due to disease (through local or systemic signaling, via soluble factors or hormones) or drug treatment.

## Results

We created a spatial, stochastic cellular automaton model for the simulation of bone remodeling, which uses the concepts of EGT for the update criteria of the simulation domain. The four states of the cellular automaton (*Resorption, Formation, Quiescence* and *Environment*) represent sites of activity of the actors of bone remodeling (osteoclasts, osteoblasts, osteocytes and environmental impacts) (Fig 1A-B). Per iteration, one voxel proliferates (spreads) to one of its neighboring voxels. The initial configuration can expand during the simulation, when the initial boundaries are exceeded by a state different from *Environment*. Furthermore, we restrict some interactions to prevent non-physical behavior (see Materials and methods). The core of the model are the payoff parameters, which encode the impact that one voxel state has on the spreading of another one and therefore represent biological interactions (Fig 1C). The payoff parameters range from −1 to 1. Positive values stand for a stimulating impact, negative values for an inhibitory impact. For example, a *Formation*-voxel turns into a *Quiescence*-voxel only if there is a *Quiescence*-voxel in its immediate neighborhood. Consequently, a *Quiescence*-voxel needs to expand into the *Formation*-voxel. In this context, the *Formation*-voxel needs to have a positive influence on the expansion rate of the *Quiescence*-voxel (i.e. *g*_*QF*_ has a positive value). For a more compact display, the payoff parameters are depicted in the payoff matrix (Fig 1D).

**Fig 1.**
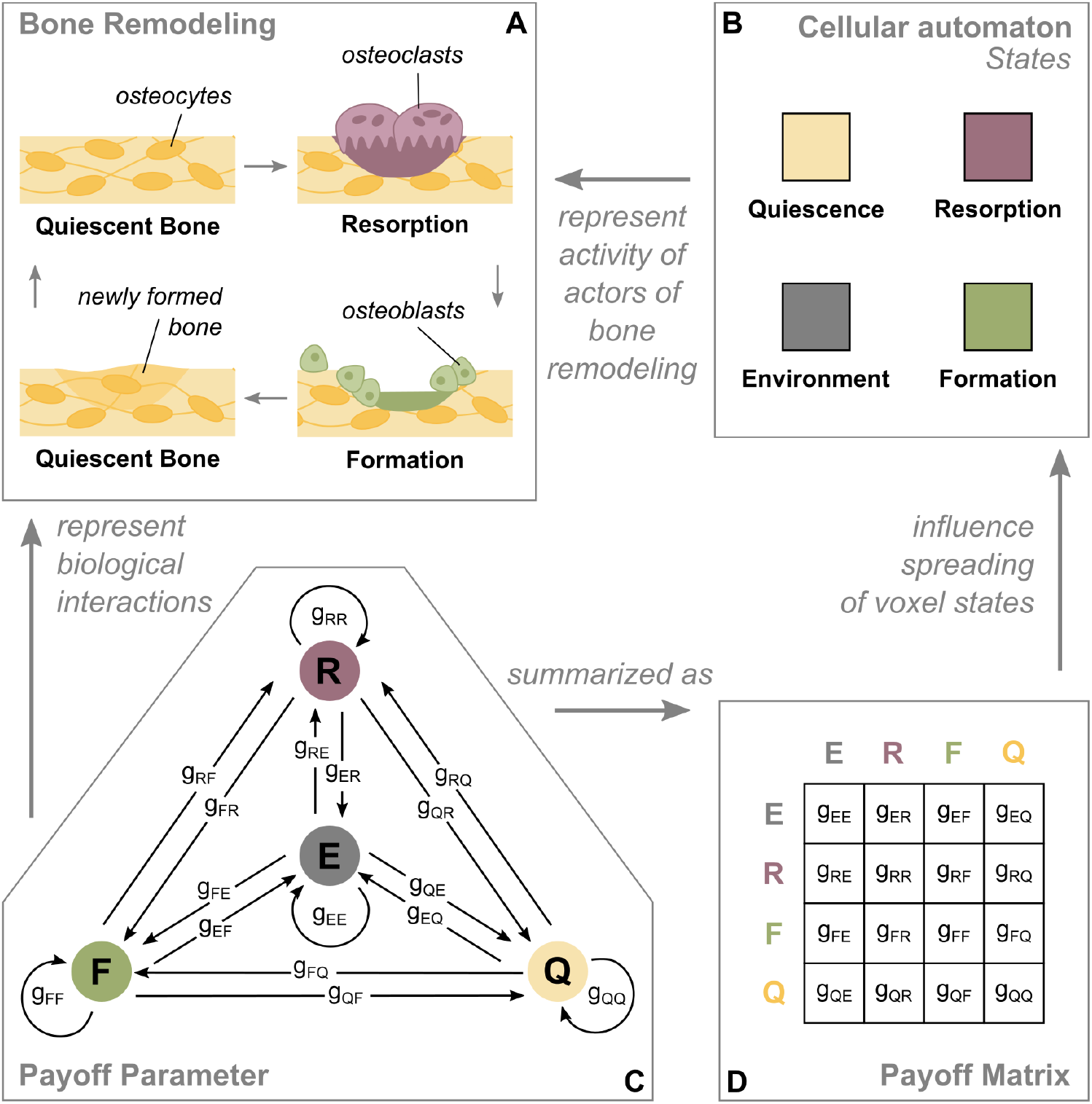
Relation of the cellular automaton model and the biology of bone remodeling. A: During bone remodeling old bone gets resorbed by osteoclasts, followed by new bone being formed by osteoblasts for the purpose of maintenance. Mechanosensitive osteocytes located in the quiescent bone mediate that process. B: The four states of the cellular automaton represent the activity of the actors of bone remodeling. C, D: The payoff parameters represent the impact that one voxel state has on the spreading of another one. They summarize all biological interactions taking place between two states and can be interpreted as stimulating (positive value) or inhibiting (negative value).

### Single patch simulations

To reduce the complexity resulting from multiple interactions, we first examine each sub-process of bone remodeling (*Resorption* and *Formation*) on its own. We set up simulations with either a *Resorption*- or a *Formation*-patch of 3×3 voxels positioned on the surface of a cube made of 28×28×28 *Quiescence*-voxels. In each case, the number of relevant payoff parameters reduces down to three (Fig 2). To address the stochastic nature of the model, each parameter combination runs for 20 rounds with a different random seed per round.

**Fig 2.**
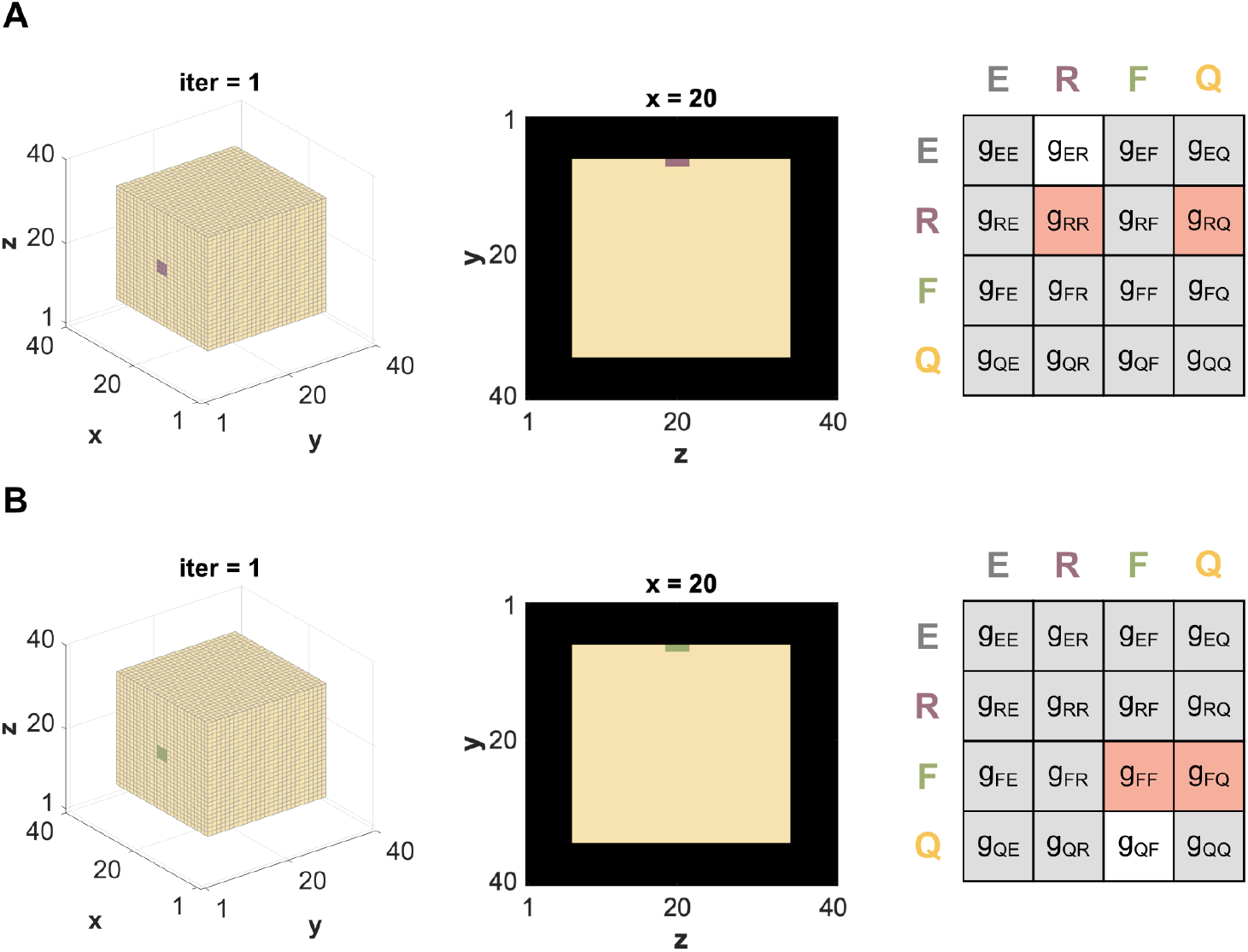
Setup for single patch simulations. A: Initial configuration for the single patch simulation of *Resorption* (lilac) on a *Quiescence*-cube, where the relevant payoff parameters are *g*_*ER*_, *g*_*RR*_ and *g*_*RQ*_. B: Initial configuration for the single patch simulation of *Formation* (green) on a *Quiescence*-cube, where the relevant payoff parameters are *g*_*QF*_, *g*_*FF*_ and *g*_*FQ*_. All parameters marked in grey are assumed to be zero. The parameters marked in white have a constant value, while the parameters marked in orange are screened from -1 to 1

#### Resorption

When simulating the resorption process of bone remodeling (Fig 1A), the main underlying interactions are between osteoclasts and the quiescent bone. During bone remodeling, resorption initiates (seeding of *Resorption*-patch), progresses for a limited amount of space and time and then stops/disappears. The main payoff parameters influencing the progression of the *Resorption*-state are *g*_*RR*_ and *g*_*RQ*_. In addition, the speed of demineralization, that is, *Environment* -voxels spreading to *Resorption*-voxels, is denoted by *g*_*ER*_ (Fig 2A).

We vary the parameters *g*_*RR*_ and *g*_*RQ*_ between −1 (maximal inhibition) and 1 (maximal stimulation) and set *g*_*ER*_ to 0.5. We group the results in a phase diagram containing three different categories: not active *Resorption* (Fig 3A, grey), bounded spreading of *Resorption* (Fig 3A, yellow to orange) and not bounded (Fig 3A, lilac). In the not active case, the *Resorption*-voxels are taken over by *Environment* -voxels, before they can spread. In the case of bounded spreading, the *Resorption*-patch starts to spread and stops within the simulation time. In the case of not bounded spreading, the *Resorption*-patch starts to spread and does not stop within the simulation time (Fig 3C, S1 Video)

**Fig 3.**
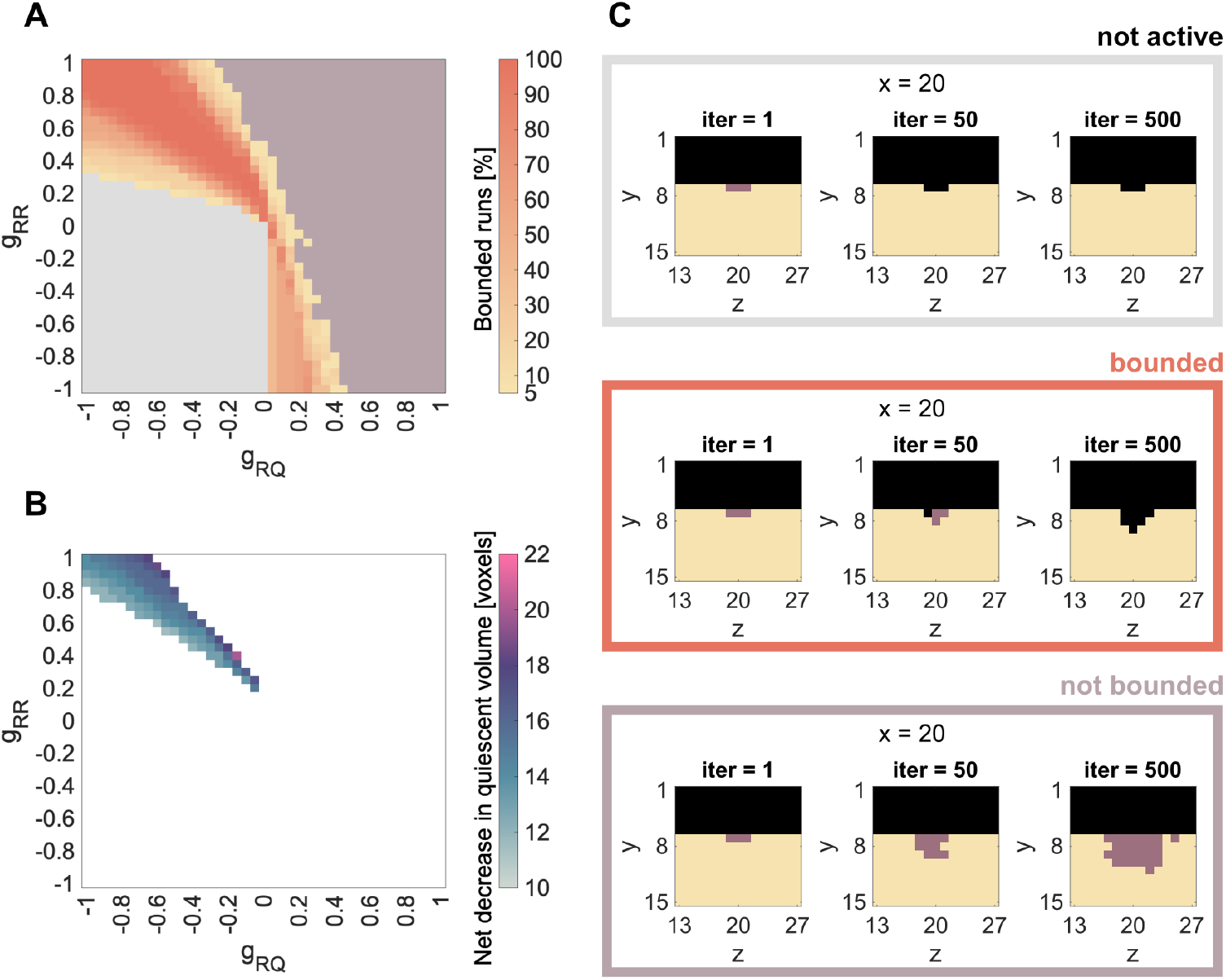
Results for single patch simulations: *Resorption*. A, C: Summary of all simulations over the combination of parameters *g*_*RR*_ and *g*_*RQ*_ and spreading activity of the seeded *Resorption*-patch. Simulation results are grouped in always not active (grey) or always not bounded spreading (lilac). Between those two phases we find parameter combinations, which lead to bounded spreading in either some (< 100%) or all (100%) of the 20 simulation runs. B: Median of final resorbed volume for parameter combinations that lead to bounded spreading in all runs.

To account for stochasticity, we run 20 independent simulations for each of the relevant parameter combinations. Some of the parameter combinations are either always, i.e. 20 out of 20, not active (grey) or always lead to not bounded spreading (lilac). Between those two phases we find parameter combinations that lead to bounded *Resorption* in either some (< 100%) or all (100%) of the 20 rounds (Fig 3A). Whenever there is a combination of two negative parameters (bottom left corner), the result is always not active, since the impact of both parameters (*g*_*RR*_ and *g*_*RQ*_) on the *Resorption* patch is inhibitory. Whenever both parameters have high positive value (meaning a strong stimulation), the spreading is not bounded anymore (upper right corner). The bounded spreading phase is located between those two extremes and displays a yellow-orange gradient indicating the percentage of bounded spreading runs out of the 20 simulations. This result remains qualitatively the same, when changing the demineralization speed *g*_*ER*_ (S1 FigA), the size of the *Resorption* patch (S2 FigA) or the position of the *Resorption*-patch (S3 FigA).

Biologically, the case of bounded spreading is the most relevant, since this is the only dynamic that can lead to an effective but limited remodeling process. Therefore, the phase within the parameter domain leading in 100% of the simulations to bounded *Resorption* is the most applicable. This phase is mainly situated in the upper left corner of the phase diagram, which represents positive *g*_*RR*_ values and negative *g*_*RQ*_ values. This can be interpreted as a stimulating impact of osteoclast signalling on osteoclasts (*g*_*RR*_ > 0) and an inhibitory impact of osteocyte signalling on osteoclasts (*g*_*RQ*_ < 0). The diagonal alignment of this phase suggests that the higher the stimulation from the osteoclasts, the higher the inhibition required from the osteocytes to keep *Resorption* bounded. Furthermore, that phase is more stable when the absolute value of both parameters is high. Focusing on only those parameter combinations leading in 100% of the simulations to bounded *Resorption*, we evaluate the median of net decrease in quiescent volume (Fig 3B). Here we find, that on the border to the not bounded spreading phase more voxels are resorbed than on the border to the not active phase.

#### Formation

When simulating the formation process of bone remodeling (Fig 1A), the main underlying interactions are between osteoblasts and the quiescent bone. During bone remodeling, formation initiates (seeding of *Formation*-patch), progresses for a limited amount of space and time and then stops/disappears. The main payoff parameters influencing the progression of the *Formation*-state are parameters *g*_*FF*_ and *g*_*FQ*_. In addition, the speed of mineralization, that is, *Quiescence*-voxels spreading to *Formation*-voxels, is denoted by *g*_*QF*_ (Fig 2B).

We vary the parameters *g*_*FF*_ and *g*_*FQ*_ between −1 and 1 and find patterns similar to the *Resorption*-simulations. Again, we group the results in a phase diagram containing three different categories: not active *Formation* (Fig 4A, grey), bounded spreading of *Formation* (Fig 4A, yellow to orange) and not bounded (Fig 4A, green). In the not active case, the *Formation*-voxels are taken over by *Quiescence* before they can spread (Fig 4C), S2 Video)

**Fig 4.**
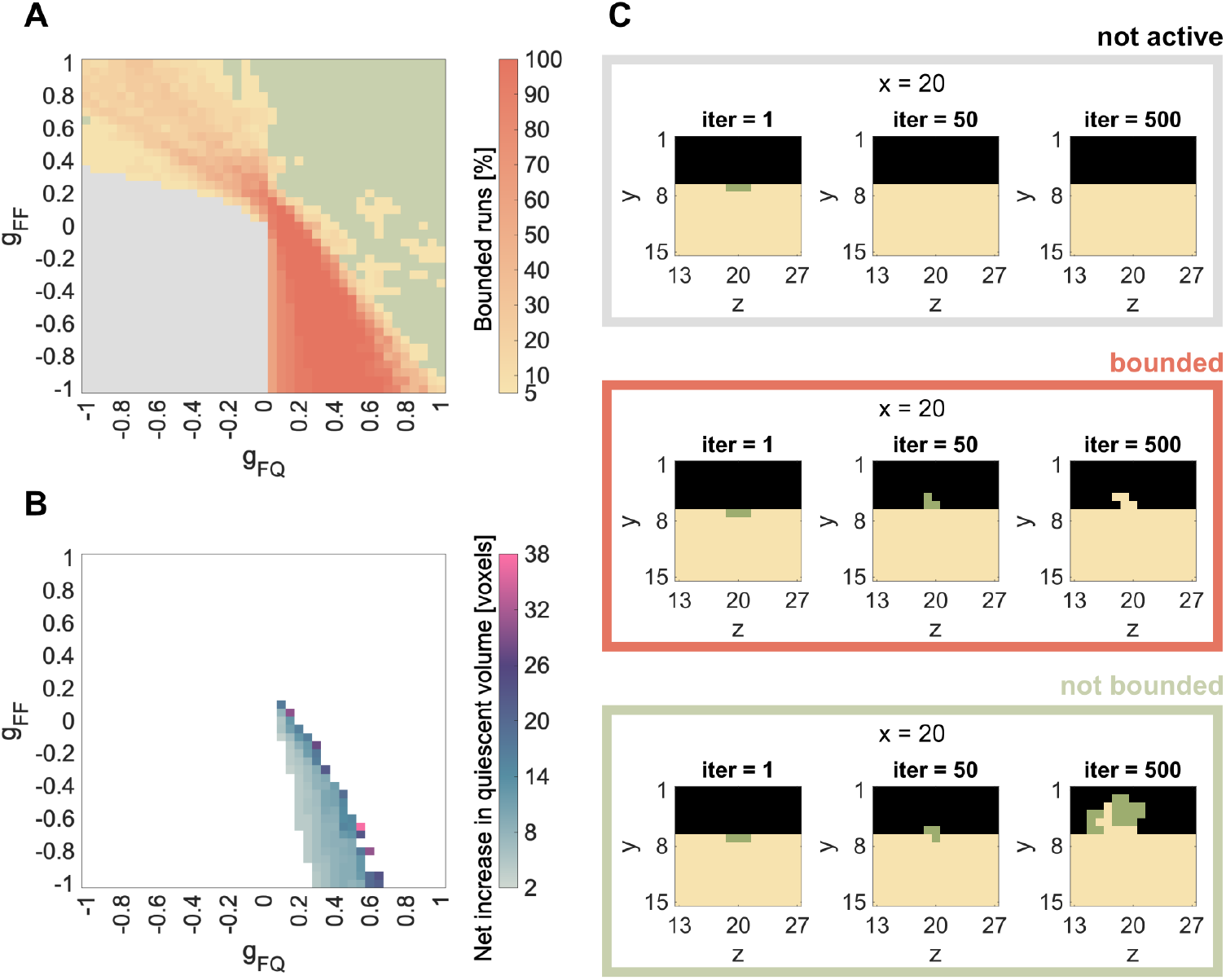
Results for single patch simulations: *Formation*. A, C: Summary of all simulations over the combination of parameters *g*_*FF*_ and *g*_*FQ*_ and spreading activity of the seeded *Formation*-patch. Simulation results are grouped in always not active (grey) or always not bounded spreading (green). Between those two phases we find parameter combinations, which lead to bounded spreading in either some (< 100%) or all (100%) of the 20 simulation runs. B: Median of final formed volume for parameter combinations that lead to bounded spreading in all runs.

Again, we run 20 independent simulations for each of the relevant parameter combinations. Some of the parameter combinations are either always, i.e. 20 out of 20, not active (grey) or always lead to not bounded spreading (green). Between those two phases, we find parameter combinations, which lead to bounded *Formation* in either some (< 100%) or all (100%) of the 20 rounds (Fig 4A). Areas where *Formation* is never active, are situated in the bottom left corner, while areas where *Formation* is never bounded, are situated in the upper right corner (Fig 4A). The bounded spreading phase is located between those two extremes and displays a yellow-orange gradient indicating the percentage of bounded spreading runs out of the 20 simulations. The three different phases also remain for *Formation*, when changing the mineralization speed *g*_*QF*_ (S1 FigB), the size of the *Formation* patch (S2 FigB) or its position (S3 FigB).

The bounded *Formation* (100% of the simulations lead to bounded spreading) and the parameter combinations leading to it are the most relevant. They are mainly situated in the bottom right corner of the phase diagram, which represents negative *g*_*FF*_ values and positive *g*_*FQ*_ values. This can be interpreted as an inhibitory impact of osteoblast signalling on osteoblasts (*g*_*FF*_ < 0) and a stimulating impact of osteocyte signalling on osteoblasts (*g*_*FQ*_ > 0). In contrast to *Resorption*, the bounded *Formation* is not clearly aligned diagonal, but rather spans through the lower left triangle of the fourth quadrant. It is more stable, for a high inhibitory impact of osteoblasts on osteoblast and gets less stable as this inhibitory impact declines. This is further discussed in the Discussion section. Focusing on only those parameter combinations that lead to 100% bounded spreading, we evaluate the median of the net increase in quiescent volume (Fig 4B). Again, we find that on the border to the not bounded spreading phase more voxels are formed than on the border to the not active phase.

### Effect of the size of the neighborhood

The choice of the cellular automaton neighborhood has a strong influence on the overall simulation dynamics. The neighborhood in the single patch simulations is initially set to a von-Neumann neighborhood (radius = 1), meaning that only voxels sharing a surface with the voxel in question are considered neighbors (Fig 5A). For a 3D grid of cubic cells this involves six voxels. To understand the role of the size of the microenvironment on the model dynamics, the single patch simulations are repeated with the next larger neighborhoods also typical for cellular automaton models. For this we choose a Moore neighborhood, meaning that all voxels sharing a surface or a corner with the voxel in question are considered neighbors. Two sizes of Moore neighborhood are considered: radius = 1 with 26 voxels on a cubic grid and radius = 2 with 124 (Fig 5B-C). Finally, a mean field approximation, where all voxels in the simulation are considered neighbors represents the largest neighborhood possible (Fig 5D).

**Fig 5.**
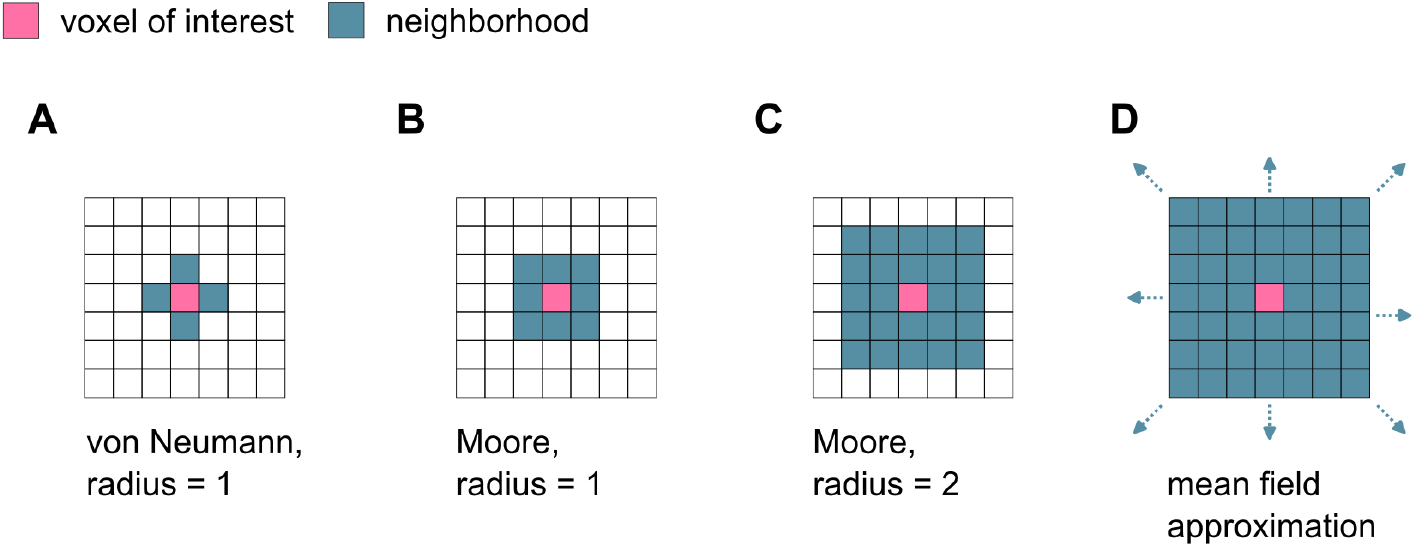
Different neighborhoods of cellular automata. A: Von-Neumann neighborhood with radius 1 (6 neighbors in 3D). B, C: Moore neighborhood with r=1 (26 neighbors in a cubic grid) or r=2 (124 neighbors in a cubic grid). D: For the mean field approximation all voxels in the simulation domain are considered neighbors.

When increasing the size of the neighborhood only slightly beyond the nearest neighbors (Moore *r* = 1, Fig 5B), the phase of bounded spreading in both cases (*Resorption* and *Formation*) becomes narrower, and tilts its position to the middle of the parameter domain (Fig 6B). Further expansion of the neighborhood (Moore *r* = 2, Fig 5C) leads to an even narrower bounded spreading phase, which is situated around *g*_*RQ*_ = 0 and *g*_*FQ*_ = 0 for *Resorption* and *Formation*, respectively (Fig 6C). This approaches the bounded spreading phase of the mean field approximation with a single vertical line (Fig 6D). Increasing the size of the neighborhood also leads to matching positions of the bounded spreading phases of *Resorption* and *Formation*.

**Fig 6.**
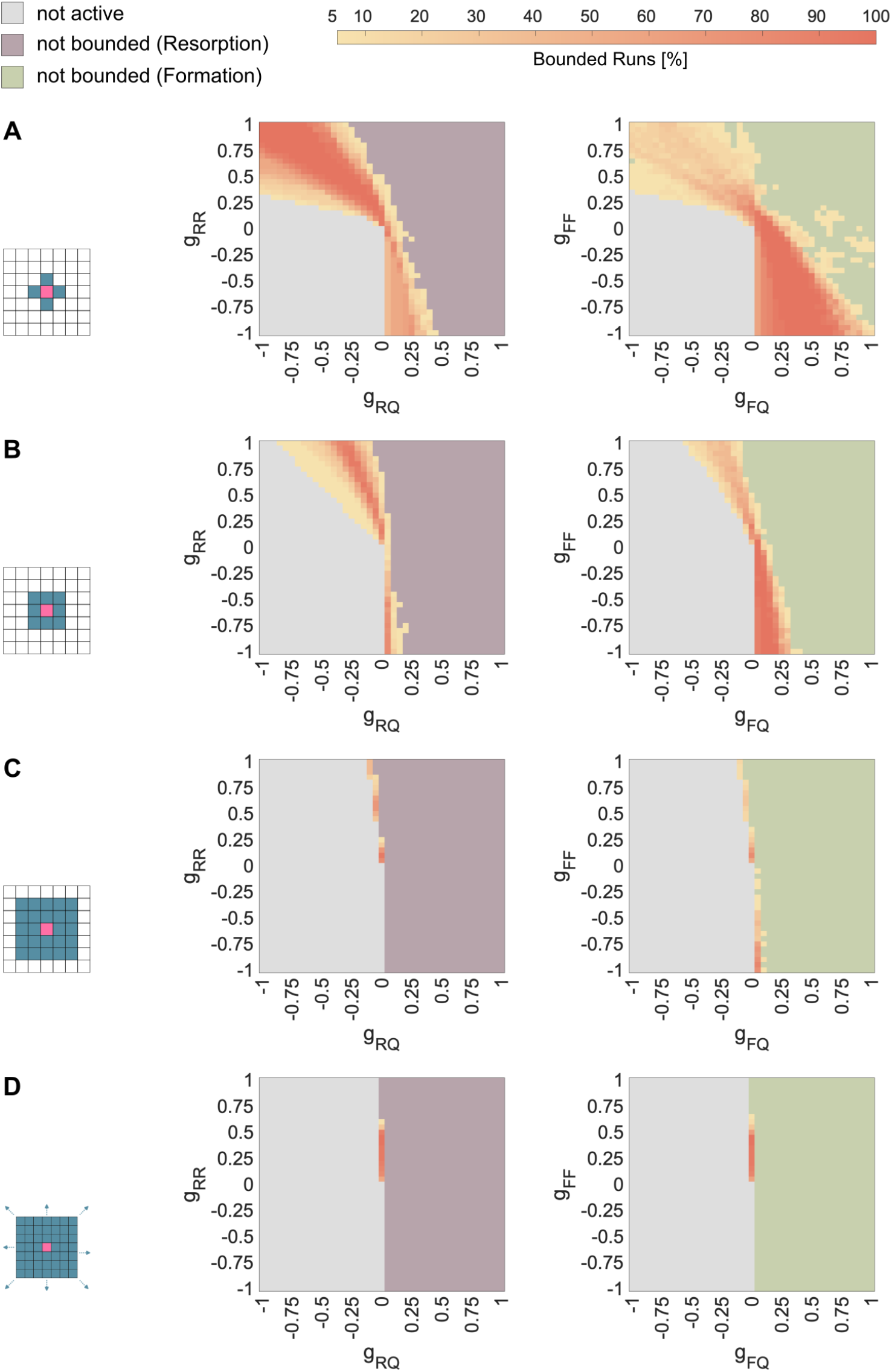
Effect of the size of the neighborhood on the bounded spreading phase of *Resorption* and *Formation*. Bounded spreading phase for *Resorption* (left) and *Formation* (right) with A: Von-Neumann neighborhood with radius=1 (6 neighbors) B: Moore neighborhood with *radius* = 1 (26 neighbors) C: Moore neighborhood with *radius* = 2 (124 neighbors) D: Mean field approximation (all voxels are neighbors)

### The remodeling cycle: Combining *Resorption* and *Formation*

To simulate one cycle of bone remodeling, we now use the results from the single patch simulations, focusing on the parameter sets that lead in 100% of rounds to bounded spreading. For this we choose different combinations of speed of demineralization (*g*_*ER*_) and speed of mineralization (*g*_*QF*_). Then we combine all parameter sets for *Formation* (*g*_*FQ*_, *g*_*FF*_) and *Resorption* (*g*_*RQ*_, *g*_*RR*_), which lead in 20/20 runs to a bounded spreading phase for the respective speed. Both processes (*Resorption* and *Formation*) are seeded one after the other, thereby assuming that the two processes are separated in time. Since bone remodeling is the process of bone maintenance and homeostasis, we monitor the net change of volume at the end of the remodeling cycle.

To imitate the pattern of the bone remodeling cycle from Fig 1A, we re-fill some of the *Environment* -voxels left behind by *Resorption* with *Formation*-voxels. An example of this is shown in Fig 7A, where we combine two parameter pairs leading to bounded spreading in *Formation* (*g*_*FF*_ = −0.85, *g*_*FQ*_ = 0.4) after *Resorption* (*g*_*RR*_ = 0.9, *g*_*RQ*_ = −0.6) for *g*_*QF*_ = 0.3 and *g*_*ER*_ = 0.8 to simulate one remodeling cycle (see also S3 Video. After simulating all combinations of parameter pairs (*g*_*RR*_, *g*_*RQ*_) and (*g*_*FF*_, *g*_*FQ*_) for the three combinations of (de-)mineralization speed (*g*_*ER*_ < *g*_*QF*_, *g*_*ER*_ = *g*_*QF*_ and *g*_*ER*_ > *g*_*QF*_), we then quantify the total volume change at the end of the remodeling cycle.

**Fig 7.**
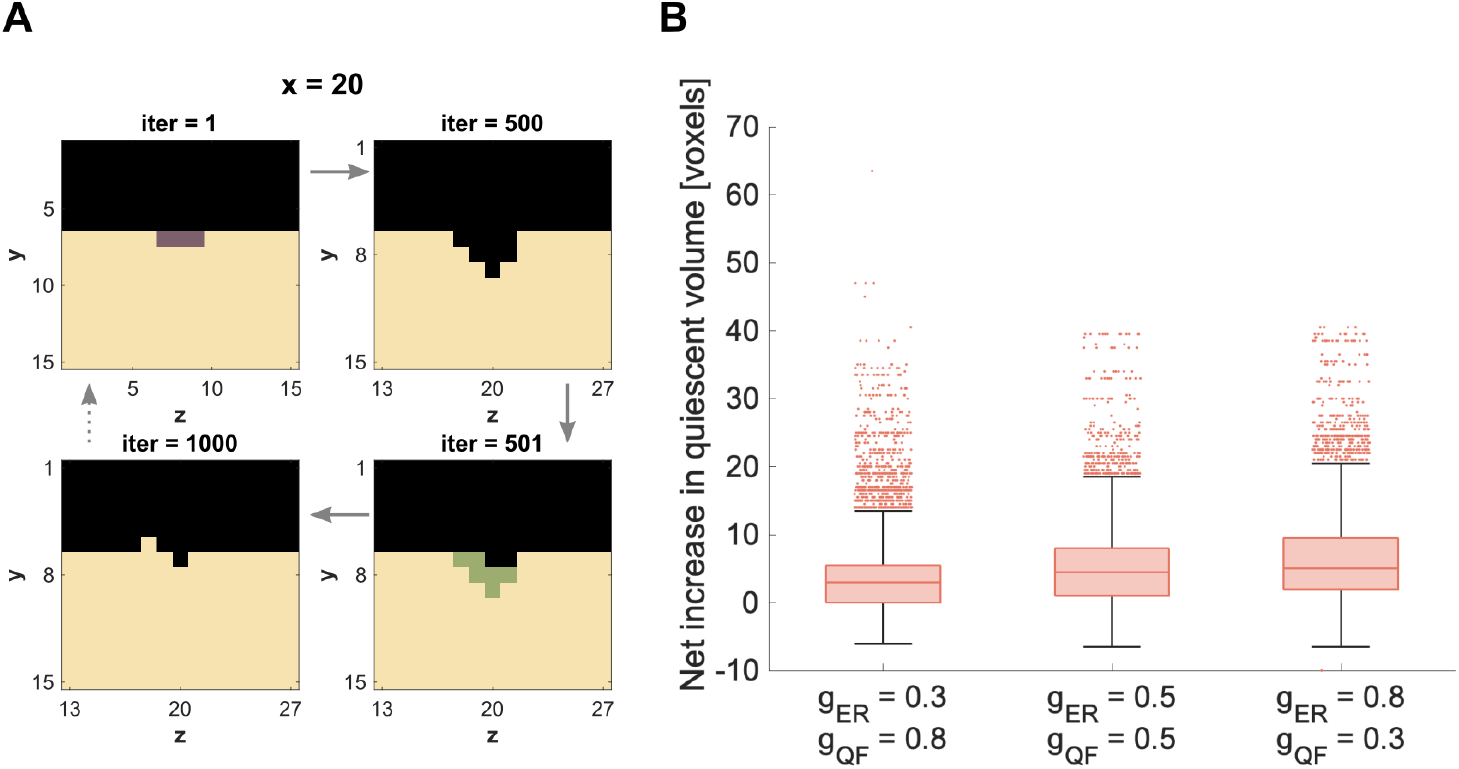
Combining bounded spreading of *Resorption* and *Formation*. A: Exemplary simulation of a bone remodeling cycle consisting of *Resorption*… (iter=1-500) followed by *Formation* (iter=500-1000). B: Distribution of the net increase in quiescent volume (median of 20 runs) over all combinations of chosen parameter sets excluding those with not bounded *Formation* (for details see S1 Table).

In contrast to the single patch simulations, some parameter combinations lead to unbounded *Formation* in individual runs. This affects less than 2.5% of total parameter combinations for each of the three cases (for details see S1 Table). We excluded all parameter combinations with unbounded *Formation* runs for the evaluation in Fig 7B. The distribution of the net increase in volume, however, differs only slightly compared to the evaluation of all parameter combinations (without exclusions). The median is the same for both evaluations (for comparison see S4 Fig).

In all three combinations of (de-)mineralization speed, the median of the total volume change is slightly larger than zero (Fig 7B). This implies a net volume growth at the end of a remodeling cycle. This tendency increases with the ratio of speed of demineralization to mineralization, 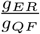.

## Discussion

With our cellular automaton model, we can explore the complex interactions necessary for a well-balanced bone remodeling cycle. Starting with simple simulations, we established a phase diagram for parameters leading to bounded spreading of *Resorption* and *Formation*, ultimately enabling the simulation of a bone remodeling cycle. The model allows us to compare the identified parameter spaces with cellular mechanisms known from literature.

### Resorption

With the phase diagram of the single patch *Resorption* simulations we showed how changing either one of the studied parameters (*g*_*RQ*_ or *g*_*RR*_) can change the whole dynamic of the resorption process: from inactive *Resorption* over bounded to unbounded *Resorption* (Fig 3A). We considered the phase of bounded *Resorption* as physiological behavior, since *Resorption* progresses first, but stops at a certain point. We further observed variation in the amount of resorbed bone within the bounded spreading phase (Fig 3B). This, again, was a result of varying one or both of the studied parameters.

When we investigate the parameter combinations leading to bounded *Resorption*, we can put this in context with current discussions on bone remodeling signaling. Biologically, *Resorption* is mainly stimulated through macrophage colony stimulating factor (MCSF) and receptor activator of nuclear factor *κ*B (RANK) originated from cells of the osteoblast lineage [35]. Those, however, should not be confused with active osteoblasts involved in bone formation, but rather include osteocytes within the bone matrix, bone lining cells or osteoblast progenitors [36] [35]. Consequently, this stimulation is represented by the parameter *g*_*RQ*_. Furthermore, parameter *g*_*RR*_ represents autocrine regulation of osteoclast formation by cytokines interleukin-1*α* (IL-1*α*) and tumor necrosis factor *α* (TNF-*α*), as well as osteoclast progenitors recruitment regulated by sphingosine-1-phosphate (S1P) derived from osteoclasts [33] [36].

When both parameters (*g*_*RQ*_ and *g*_*RR*_) are positive (therefore stimulating) the model leads to unbounded spreading of *Resorption* (Fig 3A). For bounded spreading, one of the two stimuli needs to be negative. Moreover, the bounded *Resorption* phase is more stable for a strong *g*_*RR*_ stimulus than for a strong *g*_*RQ*_ stimulus. The fact that all of the 100% bounded spreading runs are located at positive *g*_*RR*_ and negative *g*_*RQ*_ stimulus, suggests that living osteocytes negatively regulate osteoclasts [35]. According to this, the recruitment of osteoclasts is caused by osteocyte death (due to microcracks, low oestrogen levels, etc.) and the consequently weakened (meaning less negative *g*_*RQ*_) regulation of osteoclasts. Possible spikes of stimulation by osteocyte death to initiate *Resorption* (recruit osteoclasts [39], [40]) are not yet captured by the current model. Here, we simulate the initiation by seeding a patch of *Resorption* voxels in the initial configuration.

### Formation

With the phase diagram we obtained from the parameter analysis of the *Formation* single patch simulations, we showed how changing either one of the studied parameters (*g*_*FQ*_ or *g*_*FF*_) can change the whole dynamic of the formation process similar to *Resorption*. We found the same three phases: inactive, bounded and unbounded *Formation* (Fig 4A). Again, we considered the phase of bounded *Formation* as physiological behavior and observed variations in the amount of formed bone within the bounded spreading phase (Fig 4B). The differences between the shape of the bounded spreading of *Resorption* (Fig 3A) and the bounded growth of *Formation* (Fig 4A) arose from the fact that *Resorption* spread into the *Quiescence*-voxels, which had an impact on the seeded *Resorption*/*Formation*-voxels, whereas *Formation* spread into the *Environment* -voxels, which had no impact (yet). Therefore, *Formation* was more tolerant about a stimulating impact from *Quiescence* (*g*_*FQ*_) than *Resorption* (*g*_*RQ*_).

When we investigate the parameter combinations leading to bounded *Formation*, we can link this with current discussions on bone remodeling signaling. For formation, osteoblasts are stimulated and/or inhibited through bone matrix either by osteocytes (stimulation through messengers, inhibition through sclerostin) [35] or osteocalcin (inhibition) [34] and biglycan (stimulation) [34]. This is represented by the parameter *g*_*FQ*_. The parameter *g*_*FF*_ describes the impact of *Formation* on *Formation* and includes but is not limited to the release of osteoblast-derived insulin-like growth factor 1 (IGF-1), which leads to osteoblast differentiation [36]. Additional stimulation through growth factors released during demineralization of the bone matrix is not represented for now. This concerns the impact of *Environment* on *Formation* (*g*_*FE*_), but would be limited to *Environment* voxels that were recently generated due to *Resorption*. Technically, this would be possible by tracking the age of each voxel. For now, however, the purpose is to keep the model as simple as possible, which includes keeping constant values for the payoff parameters in space and time and assuming the impact of *Environment* (*g*_*iE*_) to be zero.

Similar to the results for the *Resorption* patch, the spreading of *Formation* is unbounded when both parameters (*g*_*FF*_ and *g*_*FQ*_) are stimulating (Fig 4A). Most of the 100% bounded spreading runs are located at positive *g*_*FQ*_ and negative *g*_*FF*_ stimulus, which suggests that stimulation through the bone matrix is the most important mechanism [35]. Cases of bounded spreading can also be found for positive *g*_*FF*_ and negative *g*_*FQ*_, although never for 20 runs. This suggests that, for bounded *Formation*, positive stimulation through autocrine factors can to be compensated by negative regulation through the bone matrix, but this regulation would not be stable. To our knowledge autocrine inhibition (therefore *g*_*FF*_ < 0) of osteoblasts is not discussed in the literature, yet.

### Effect of the size of neighborhood

Comparison of different neighborhood sizes demonstrated that spatial modeling with nearest neighbors can reveal local control mechanisms of the microenvironment that are otherwise lost in the mean field approximation (Fig 6). This agreed with previous findings of [29], who showed for certain parameters spaces that an unstable EGT dynamic can stabilize when switching from a non-spatial to a spatial model.

The size of the bounded *Resorption* phase became narrower with increasing size of the neighborhood (Fig 6). The same applied for *Formation* (Fig 6). Furthermore, comparing neighborhood sizes showed, that differences between the shape of bounded *Resorption* (Fig 3A) and bounded *Formation* (Fig 4A) in a 6-voxel-neighborhood reduced progressively with an increasing neighborhood. This confirmed, that those differences arose from different voxel-types, and therefore directions, in which *Resorption* and *Formation* spread: the former into the *Quiescence*-voxels, the latter into the *Environment* -voxels. This showed how regulation mechanisms that work locally within a small microenvironment can be overlooked, when modeling with too large a neighborhood or a mean field approximation.

### The remodeling cycle: Combining *Resorption* and *Formation*

The combination of bounded-spreading *Resorption* and *Formation* gave promising first results of simulating a whole bone remodeling cycle using parameters from the bounded spreading phase. Quantitatively the results were still tilted towards bone growth instead of preserving the bone volume (maintenance). This suggested that if the processes of *Formation* and *Resorption* would be repeated over and over again (simulating a constant remodeling behaviour), the simulated domain would most likely increase its volume over time. This however does not mirror the maintaining nature of the bone remodeling process.

The fact that some parameter combinations for *Formation* always led to bounded spreading in the single patch simulations, but not when combining *Resorption* and *Formation*, emphasized again the importance of the local environment. When simulating the remodeling cycle, *Resorption* acted as anticipated from the results of the single patch simulations, because it was seeded in the same way (3×3 voxel-patch on the flat surface). *Formation*, however, was seeded differently: Instead of seeding on a flat surface, we based the spatial distribution of the *Formation* voxels on the *Environment* -pattern that *Resorption* left behind. This caused in some cases differences in behavior of *Formation* compared to the single patch simulations.

### Perspective

In Fig 3B and Fig 4B we show that different parameter combinations within the phase of bounded spreading lead to different amounts of net decrease and increase in quiescent volume, respectively. To ensure maintenance of the volume, the parameter pairs for *Resorption* (*g*_*RR*_ and *g*_*RQ*_) and *Formation* (*g*_*FF*_ and *g*_*FQ*_) might need to be adjusted to each other. This could lead to a sub-phase within bounded spreading, that results in not just bounded but in physiological bone remodeling. Alternatively, some of these parameters (e.g. *g*_*FF*_ and *g*_*FQ*_) could be dependant on the values of others (e.g. *g*_*RR*_ and *g*_*RQ*_).

Criteria for distinguishing physiological bone remodeling from pathological bone remodeling may be deduced from the experimental investigations of Young and colleagues [41], [42]. The spatially explicit nature of our model enables the integration of bone geometry, as characterized in the experimental data set, to serve as the initial configuration for the model. This should allow for a direct juxtaposition of the bone remodeling patterns observed experimentally with those forecasted by the model. Nevertheless, to effectively implement this approach, a constellation of technical and theoretical challenges must be addressed: the temporal dimension of the simulations must be harmonized with the biological tempo of bone remodeling by calibrating the rates within the pay-off matrix; the biological variation between individual mice requires an evaluation of the margin of error within which the model reliably mirrors the experimental findings; and, while the experimental data captures alterations in quiescent bone, the model simulates voxel dynamics over time. We remain optimistic that forthcoming research endeavors will resolve these complexities.

So far, the impact of the *Environment* state is kept neutral (without any impact) and consequently the parameters *g*_*iE*_ are set to zero. However, systemic regulation by hormones and local regulation via soluble factors modulate bone physiology. Alterations of these as well as the effect of drugs could be intergrated and modelled through the *Environment* -state. Concretely this means to investigate the influence of the parameters *g*_*iE*_ on the previously discussed phase diagrams.

For now, the number of *Formation* voxels being seeded after resorption is the same as the previously seeded *Resorption* voxels. The legitimization of that assumption can be further investigated by varying the number of seeded *Formation* voxels and possibly coupling that number to the size of the cavity left behind by *Resorption* (Fig 7A, upper right corner).

Since both, *Resorption* and *Formation* patch, are seeded on different timescales, the impact that *Resorption* and *Formation* have on each other (*g*_*RF*_ and *g*_*FR*_) is not captured in the current setup. This assumption is supported by Sims and Martin [36] who showed that coupling between active osteoclasts and osteoblasts via direct contact is unlikely. Findings of [43], however, introduce a mixed reversal-resorption phase, where at the end of a resorption phase the density of osteoprogenitor cells grows until a certain threshold is reached and formation starts. This dynamic could be modeled by seeding the *Formation* patch while *Resorption* voxels are still present, thereby simulating the switch from reversal-resorption- to formation-phase by *Formation* voxels out competing remaining *Resorption* voxels.

In the proposed model, the *Resorption* and *Formation* patch are seeded manually. In the future, the model could be coupled with a finite element analysis similar to [12]. The resulting strains would inform the seeding of the remodeling patches. According to this, the *Resorption* patches would be seeded randomly assuming regular appearing microcracks and other damages in the bone. The *Formation* patch, however, would be seeded in areas of high strains, which can either be caused by increased load conditions or a resulting resorption cavity. Furthermore, the magnitude of strain could also serve as a criterion for the number of *Formation* voxels being seeded.

## Conclusion

With the proposed model we laid the foundations for simulating spatial bone remodeling with a stochastic cellular automaton using evolutionary game theory. Such modeling approach allowed us generate phase diagrams of bone resorption and formation to qualitatively understand the impact and sensitivity of stimulatory/inhibitory effects of certain biological processes.. Furthermore, comparing the effect of different neighborhood sizes demonstrated that spatial modeling with nearest neighbors can reveal local control mechanisms of the microenvironment that are lost in the mean field approximation.

In the future, the proposed model could inform *in vitro*/*in vivo* experimental design and/or be combined with other *in silico* models investigating external impacts on bone remodeling (finite element analysis of mechanical loading, cancer models or drug/treatment protocols). To sum up, this model opens the window to explore the complex interplay of factors essential for healthy bone remodeling and potential alterations due to disease (through local or systemic signaling, via soluble factors or hormones) or drug treatment.

## Materials and methods

### Model

A spatial four strategy model resembling a stochastic cellular automaton, whose update rule is based on criteria borrowed from EGT is set up in Matlab (version 2023b). The simulation domain is approximated by a cube and discretized into n same-sized-voxels per axis. Each voxel gets assigned a voxel state. Based on the process of bone remodeling, the voxel states can vary between *Environment* (= 1), *Resorption* (= 2), *Formation* (= 3) and *Quiescence* (= 4) (Fig 1A-B). The current state of a voxel with position **x** is described by *ξ*(**x**) ∈ {1, 2, 3, 4}. We adopt the notation of Ryser et al [29], where the payoff parameter *g*_*ij*_ describes the impact each state *j* has on the spreading rate of the state *i* (Fig 1C). The set of voxels previously defined as neighbors of voxel **x** is denoted by **y** ∼ **x**. The neighborhood is set to include only the nearest neighbors, meaning those which share a surface with the voxel of interest (Fig 5A).

All interactions described with payoff parameters *g*_*ij*_ are coded in a payoff matrix (Fig 1D), which contains sixteen entries for a system with four strategies. They are decisive for the dynamics of the whole system. The payoff parameters range between -1 (maximally inhibitory) and 1 (maximally stimulating) to capture their relative proportion to each other. The payoff parameters are constant in space and time and are defined at the beginning of the simulation.

The dynamics of the simulated system are calculated by applying a Gillespie algorithm, which uses the expansion rates of the voxels to determine their probability to spread. The final expansion rate for each voxel per iteration is calculated by summing up the impact from all neighboring voxels **y**:

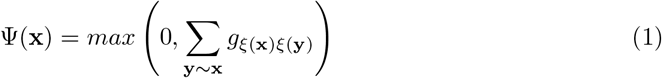

After the spreading rates Ψ(**x**) for all voxels have been calculated, each voxel **x** gets assigned a random time exponentially distributed with mean time Ψ(**x**)^−1^ (with Ψ(**x**)^−1^ = ∞ if Ψ(**x**) = 0). The voxel that gets assigned the shortest time spreads into one of the k voxels within the neighborhood, chosen uniformly at random.

Before spreading, the neighborhood states of this voxel are checked for non-physical changes (Table 1), which are then excluded (e.g. *Formation* spreading to *Quiescence* or *Quiescence* spreading to *Resorption*). The decision whether the spreading from *Resorption* to *Formation* and vice versa are to be excluded or not is up for discussion and needs further investigation. For this work, we model *Resorption* and *Formation* on separated time scales. Consequently, they are not simultaneously present in the model domain and our results are independent of that decision.

**Table 1.** Allowed spreading directions of voxel states.

| spreading state | Environment | Resorption | Formation | Quiescence |
| --- | --- | --- | --- | --- |
| allowed spreading to | Environment<br>Resorption | Resorption<br>Quiescence<br>(Formation) | Formation<br>Environment<br>(Resorption) | Quiescence<br>Formation |

As boundary condition, we assume *Environment* -voxels outside of the simulated domain which partially constitute the neighborhood of the boundary voxels. Each time the initial boundaries are exceeded by a state different from *Environment*, the domain expands by one layer of voxels on each side of the cube. This enables a growing and shrinking simulation domain.

### Single patch simulations

Due to the high number of payoff parameters, it is difficult to investigate all of them at the same time. We assume the impact of the *Environment* -voxels to be zero for this work (all parameters *g*_*iE*_ = 0, yet in the future this could include the impact of disease and drugs. Furthermore, we do not expect any interaction connected to spreading between *Quiescence* and *Environment* (*g*_*EQ*_ = 0, *g*_*QE*_ = 0).

To reduce the parameters further we set up simulations of only *Resorption* or only *Formation*, i.e. single patch simulations. For this, we seed a 3×3 voxel *Resorption*- or *Formation*-patch on the surface of a 28×28×28 voxel *Quiescence*-cube, which is surrounded by one layer of *Environment* -voxels. The whole domain has a size of 30×30×30 voxels.

In each case, this reduces the number of relevant parameters down to three. First, when simulating the single *Resorption*-patch, no *Formation* is present or can emerge and therefore all parameters *g*_*iF*_ and *g*_*Fi*_ are zero. Also, without *Formation* present, *Quiescence* will not be able to spread beyond its initial voxels so we can assume parameter *g*_*QR*_ as zero as well. Second, the argumentation line for the single *Formation*-patch simulation is similar. All parameters *g*_*iR*_ and *g*_*Ri*_ are zero. Also, without *Resorption* present, *Environment* will not be able to spread beyond its initial voxels so we can assume parameter *g*_*EF*_ as zero as well.

In the end, the single *Resorption*-patch simulation is modeled with the payoff parameters *g*_*RR*_, *g*_*RQ*_ and *g*_*ER*_ and the single *Formation*-patch simulation is modeled with the payoff parameters *g*_*FF*_, *g*_*FQ*_ and *g*_*QF*_.

The parameter *g*_*ER*_ represents the rate of demineralization, does not have a direct impact on the spreading of *Resorption* and is assigned a fixed value of 0.5 (effect of values 0.3 and 0.8 in S1 FigA). Analogously, the parameter *g*_*QF*_ represents the rate of mineralization, does not have a direct impact on the spreading of *Formation* and is assigned a fixed value of 0.5 (effect of values 0.3 and 0.8 in S1 FigB). Finally, we vary *g*_*RR*_ and *g*_*RQ*_ between -1 and 1 in the single *Resorption*-patch simulation, while we vary *g*_*FF*_ and *g*_*FQ*_ between -1 and 1 in the single *Formation*-patch simulation.

Considering the stochasticity of the model, we run 20 simulations (with 20 different random seeds) for each parameter combination. Each simulation runs for 1000 iterations. We expect some occasional changes (unbounded runs becoming bounded) for a significantly larger amount of iterations. However, those occasional cases do not change the main resulting phase diagram and therefore have no influence on the conclusion.

The results are categorized in the following way: Simulations, where the patch has disappeared without initial spreading are considered not active. When the patch has spread, but still disappeared at the end of the simulation, it is considered as bounded spreading. Simulations, where the patch has not disappeared by the end of the 1000 iterations, are considered as unbounded spreading. Depending on the parameter combination, all 20 simulation runs can either fall in the same category or can fluctuate between different ones.

### Effect of the size of neighborhood

To investigate the impact of the size of the neighborhood on the model dynamics, we repeat the single patch simulations (20 runs á 1000 iterations with *g*_*ER*_ = 0.5 and *g*_*QF*_ = 0.5) for four different neighborhood sizes: von-Neumann neighborhood with radius 1 (6 neighbors in 3D), Moore neighborhood with r=1 (26 neighbors in 3D) and r=2 (124 neighbors in 3D)and mean field approximation, where all voxels in the simulation domain are considered neighbors (Fig 5). The neighborhood specifies a) which neighboring voxels are considered for the calculation of the spreading rates and b) where a voxel can theoretically spread to.

In the case of the mean field approximation, each simulation is run for one million iterations. Furthermore, runs are only considered bounded, if less than 10% of the *Quiescence*-volume have been resorbed or formed.

### The remodeling cycle: Combining *Resorption* and *Formation*

To simulate one whole bone remodeling cycle, one single *Resorption*-patch simulation is followed by one single *Formation*-patch simulation. We only use parameter combinations (*g*_*RR*_ and *g*_*RQ*_; *g*_*FF*_ and *g*_*FQ*_), which lead to a bounded spreading in all runs of the single patch simulations. To investigate the relation of *g*_*ER*_ and *g*_*QF*_, we simulate three different combinations: *g*_*ER*_ = 0.3 and *g*_*QF*_ = 0.8, *g*_*ER*_ = 0.5 and *g*_*QF*_ = 0.5, and *g*_*ER*_ = 0.8 and *g*_*QF*_ = 0.3. In all three cases we combine all parameters sets that lead to bounded *Resorption* (*g*_*RR*_ and *g*_*RQ*_) and bounded for *Formation* (*g*_*FF*_ and *g*_*FQ*_). Again, we run 20 simulations (with 20 different random seeds) for each parameter combination.

The two processes of *Resorption* and *Formation* are combined on separated timescales. First, a *Resorption* patch of 3×3 voxel is set on the surface of a 28×28×28 *quiescent* cube (identical to the single patch simulation, Fig 2A). Since we only take the parameter combinations for bounded spreading, we can be sure that after 1000 iterations all *Resorption* voxels will have disappeared from the model domain. Next, we set nine *Formation* voxels in the deepest points of the cavity that *Resorption* has left behind (Fig 7A, lower right). Due to the changed *Formation* setup compared to the single patch simulation we can not expect that all *Formation*-voxels will have disappeared after 1000 iterations. Therefore, the simulation runs again until all *Formation* voxels have disappeared or the model domain exceeds a side length of 46 voxels. In case, the simulation run stops because of the latter, we exclude the respective parameter combinations (all 20 runs) from the evaluation (for details see S1 Table).

## Code availability

The source code for the introduced model is available on a GitHub repository: https://github.com/CipitriaLab/3D_CA_BoneRemodeling.

Additionally, all code written for this publication (including help functions, input files, model code and evaluation scripts) are available on Edmond: https://doi.org/10.17617/3.FNXOA3

## Supporting information

**S1 Fig. Single patch simulations varying demineralization rate** *g*_*ER*_ **and mineralization rate** *g*_*QF*_. A: Bounded spreading phase for *Resorption* with *g*_*ER*_ = 0.3 (left) and *g*_*ER*_ = 0.8 (right) B: Bounded spreading phase for *Formation* with *g*_*QF*_ = 0.3 (left) and *g*_*QF*_ = 0.8 (right).

**S2 Fig. Single patch simulations varying the size of the *Resorption* /*Formation* patch**. A: Bounded spreading phase for *Resorption* with a 5×5 voxel patch (left) and 7×7 voxel patch (right) B: Bounded spreading phase for *Formation* with a 5×5 voxel patch (left) and 7×7 voxel patch (right).

**S3 Fig. Single patch simulations varying the position (depth) of the *Resorption* /*Formation* patch (dotted line)** A: Bounded spreading phase for *Resorption* with the patch positioned one, two and three layers below surface (from top to bottom) B: Bounded spreading phase for *Formation* with the patch positioned one, two and three layers below surface (from top to bottom).

**S4 Fig. Combining bounded spreading of *Resorption* and *Formation*** A: Distribution of net increase of volume for all parameter combinations (without exclusion) B: Distribution of net increase of volume excluding parameter combinations of unbounded runs.

**S1 Video. Time sequence of the different categories of *Resorption* spreading shown in Fig 3C**

**S2 Video. Time sequence of the different categories of *Formation* spreading shown in Fig 4C**

**S3 Video. Time sequence of combining *Resorption* and *Formation* for one remodeling cycle shown in Fig 7A**

**S1 Table. Details for the combined simulations shown in Fig 7B**

## Acknowledgments

All simulations were carried out at the Max Planck Computing and Data Facility (MPCDF) and the computing cluster of the Max Planck Institute of Colloids and Interfaces (MPICI). A. Heller thanks the International Max Planck Research School (IMPRS) on Multiscale Bio-Systems for financial support. A. Cipitria acknowledges the funding by the Deutsche Forschungsgemeinschaft (DFG) Emmy Noether Grant CI 203/2-1 and the IKERBASQUE Basque Foundation for Science. We thank open access funding provided by the Max Planck Society. We thank Peter Fratzl, Richard Weinkamer and Sarah Young for fruitful discussions.

## References

1. Carter DR, Fyhrie DP, Whalen RT. Trabecular bone density and loading history: Regulation of connective tissue biology by mechanical energy. Journal of Biomechanics. 1987;20(8):785–794. doi:10.1016/0021-9290(87)90058-3.

2. Huiskes R, Weinans H, Grootenboer HJ, Dalstra M, Fudala B, Slooff TJ. Adaptive bone-remodeling theory applied to prosthetic-design analysis. Journal of Biomechanics. 1987;20(11-12):1135–1150. doi:10.1016/0021-9290(87)90030-3.

3. Weinans H, Huiskes R, Grootenboer HJ. The behavior of adaptive bone-remodeling simulation models. Journal of Biomechanics. 1992;25(12):1425–1441. doi:10.1016/0021-9290(92)90056-7.

4. Mullender MG, Huiskes R, Weinans H. A physiological approach to the simulation of bone remodeling as a self-organizational control process. Journal of Biomechanics. 1994;27(11):1389–1394. doi:10.1016/0021-9290(94)90049-3.

5. Prendergast PJ, Taylor D. Prediction of bone adaptation using damage accumulation. Journal of Biomechanics. 1994;27(8):1067–1076. doi:10.1016/0021-9290(94)90223-2.

6. Adachi T, Tomita Y, Sakaue H, Tanaka M. Simulation of Trabecular Surface Remodeling based on Local Stress Nonuniformity. JSME International Journal Series C. 1997;40(4):782–792. doi:10.1299/jsmec.40.782.

7. Huiskes R, Ruimerman R, van Lenthe GH, Janssen JD. Effects of mechanical forces on maintenance and adaptation of form in trabecular bone. Nature. 2000;405(6787):704–706. doi:10.1038/35015116.

8. Tayyar S, Weinhold PS, Butler RA, Woodard JC, Zardiackas LD, St John KR, et al. Computer simulation of trabecular remodeling using a simplified structural model. Bone. 1999;25(6):733–739. doi:10.1016/S8756-3282(99)00218-5.

9. Adachi T, Tsubota Ki, Tomita Y, Hollister SJ. Trabecular Surface Remodeling Simulation for Cancellous Bone Using Microstructural Voxel Finite Element Models. Journal of Biomechanical Engineering. 2001;123(5):403–409. doi:10.1115/1.1392315.

10. Guo XE, Kim CH. Mechanical consequence of trabecular bone loss and its treatment: a three-dimensional model simulation. Bone. 2002;30(2):404–411. doi:10.1016/S8756-3282(01)00673-1.

11. Weinkamer R, Hartmann MA, Brechet Y, Fratzl P. Stochastic Lattice Model for Bone Remodeling and Aging. Physical Review Letters. 2004;93(22):228102. doi:10.1103/PhysRevLett.93.228102.

12. Ruimerman R, Hilbers P, van Rietbergen B, Huiskes R. A theoretical framework for strain-related trabecular bone maintenance and adaptation. Journal of Biomechanics. 2005;38(4):931–941. doi:10.1016/j.jbiomech.2004.03.037.

13. Penninger CL, Patel NM, Niebur GL, Tovar A, Renaud JE. A fully anisotropic hierarchical hybrid cellular automaton algorithm to simulate bone remodeling. Mechanics Research Communications. 2008;35(1-2):32–42. doi:10.1016/j.mechrescom.2007.10.007.

14. Tsubota Ki, Suzuki Y, Yamada T, Hojo M, Makinouchi A, Adachi T. Computer simulation of trabecular remodeling in human proximal femur using large-scale voxel FE models: Approach to understanding Wolff’s law. Journal of Biomechanics. 2009;42(8):1088–1094. doi:10.1016/j.jbiomech.2009.02.030.

15. Van Der Linden JC, Verhaar JAN, Weinans H. A Three-Dimensional Simulation of Age-Related Remodeling in Trabecular Bone. Journal of Bone and Mineral Research. 2001;16(4):688–696. doi:10.1359/jbmr.2001.16.4.688.

16. Müller R. Long-term prediction of three-dimensional bone architecture in simulations of pre-, peri-and post-menopausal microstructural bone remodeling. Osteoporosis International. 2005;16(S02):S25–S35. doi:10.1007/s00198-004-1701-7.

17. Dunlop JWC, Hartmann MA, Bréchet YJ, Fratzl P, Weinkamer R. New Suggestions for the Mechanical Control of Bone Remodeling. Calcified Tissue International. 2009;85(1):45–54. doi:10.1007/s00223-009-9242-x.

18. Schulte FA, Zwahlen A, Lambers FM, Kuhn G, Ruffoni D, Betts D, et al. Strain-adaptive in silico modeling of bone adaptation — A computer simulation validated by in vivo micro-computed tomography data. Bone. 2013;52(1):485–492. doi:10.1016/j.bone.2012.09.008.

19. Walle M, Marques FC, Ohs N, Blauth M, Müller R, Collins CJ. Bone Mechanoregulation Allows Subject-Specific Load Estimation Based on Time-Lapsed Micro-CT and HR-pQCT in Vivo. Frontiers in Bioengineering and Biotechnology. 2021;9(June). doi:10.3389/fbioe.2021.677985.

20. Komarova SV, Smith RJ, Dixon SJ, Sims SM, Wahl LM. Mathematical model predicts a critical role for osteoclast autocrine regulation in the control of bone remodeling. Bone. 2003;33(2):206–215. doi:10.1016/S8756-3282(03)00157-1.

21. Lemaire V, Tobin FL, Greller LD, Cho CR, Suva LJ. Modeling the interactions between osteoblast and osteoclast activities in bone remodeling. Journal of Theoretical Biology. 2004;229(3):293–309. doi:10.1016/j.jtbi.2004.03.023.

22. Pivonka P, Zimak J, Smith DW, Gardiner BS, Dunstan CR, Sims NA, et al. Model structure and control of bone remodeling: A theoretical study. Bone. 2008;43(2):249–263. doi:10.1016/j.bone.2008.03.025.

23. Pivonka P, Komarova SV. Mathematical modeling in bone biology: From intracellular signaling to tissue mechanics. Bone. 2010;47(2):181–189. doi:10.1016/j.bone.2010.04.601.

24. Buenzli PR, Pivonka P, Gardiner BS, Smith DW. Modelling the anabolic response of bone using a cell population model. Journal of Theoretical Biology. 2012;307:42–52. doi:10.1016/j.jtbi.2012.04.019.

25. Ait Oumghar I, Barkaoui A, Chabrand P, Ghazi AE, Jeanneau C, Guenoun D, et al. Experimental-based mechanobiological modeling of the anabolic and catabolic effects of breast cancer on bone remodeling. Biomechanics and Modeling in Mechanobiology. 2022;21(6):1841–1856. doi:10.1007/s10237-022-01623-z.

26. Ryser MD, Nigam N, Komarova SV. Mathematical Modeling of Spatio-Temporal Dynamics of a Single Bone Multicellular Unit. Journal of Bone and Mineral Research. 2009;24(5):860–870. doi:10.1359/jbmr.081229.

27. Buenzli PR, Jeon J, Pivonka P, Smith DW, Cummings PT. Investigation of bone resorption within a cortical basic multicellular unit using a lattice-based computational model. Bone. 2012;50(1):378–389. doi:10.1016/j.bone.2011.10.021.

28. Ryser MD, Qu Y, Komarova SV. Osteoprotegerin in Bone Metastases: Mathematical Solution to the Puzzle. PLoS Computational Biology. 2012;8(10):e1002703. doi:10.1371/journal.pcbi.1002703.

29. Ryser MD, Murgas KA. Bone remodeling as a spatial evolutionary game. Journal of Theoretical Biology. 2017;418(January):16–26. doi:10.1016/j.jtbi.2017.01.021.

30. Scheiner S, Pivonka P, Hellmich C. Coupling systems biology with multiscale mechanics, for computer simulations of bone remodeling. Computer Methods in Applied Mechanics and Engineering. 2013;254:181–196. doi:10.1016/j.cma.2012.10.015.

31. Kameo Y, Miya Y, Hayashi M, Nakashima T, Adachi T. In silico experiments of bone remodeling explore metabolic diseases and their drug treatment. Science Advances. 2020;6(10):1–11. doi:10.1126/sciadv.aax0938.

32. Boaretti D, Marques FC, Ledoux C, Singh A, Kendall JJ, Wehrle E, et al. Trabecular bone remodeling in the aging mouse: A micro-multiphysics agent-based in silico model using single-cell mechanomics. Frontiers in Bioengineering and Biotechnology. 2023;11:1091294. doi:10.3389/fbioe.2023.1091294.

33. Tani-Ishii N, Tsunoda A, Teranaka T, Umemoto T. Autocrine Regulation of Osteoclast Formation and Bone Resorption by IL-1α and TNFα. Journal of Dental Research. 1999;78(10):1617–1623. doi:10.1177/00220345990780100601.

34. Hadjidakis DJ, Androulakis II. Bone Remodeling. Annals of the New York Academy of Sciences. 2006;1092(1):385–396. doi:10.1196/annals.1365.035.

35. Crockett JC, Rogers MJ, Coxon FP, Hocking LJ, Helfrich MH. Bone remodelling at a glance. Journal of Cell Science. 2011;124(7):991–998. doi:10.1242/jcs.063032.

36. Sims NA, Martin TJ. Coupling the activities of bone formation and resorption: a multitude of signals within the basic multicellular unit. BoneKEy Reports. 2014;3(September 2013):1–10. doi:10.1038/bonekey.2013.215.

37. Miller AK, Bishop RT, Li T, Shain KH, Nerlakanti N, Lynch CC, et al. The bone ecosystem facilitates multiple myeloma relapse and the evolution of heterogeneous proteasome inhibitor resistant disease. bioRxiv. 2022; p. 1–35. doi:10.1101/2022.11.13.516335.

38. Nowak MA. Evolutionary Dynamics: Exploring the Equations of Life. Havard University Press; 2006.

39. Komori T. Cell Death in Chondrocytes, Osteoblasts, and Osteocytes. International Journal of Molecular Sciences. 2016;17(12):2045. doi:10.3390/ijms17122045.

40. Andreev D, Liu M, Weidner D, Kachler K, Faas M, Grüneboom A, et al. Osteocyte necrosis triggers osteoclast-mediated bone loss through macrophage-inducible C-type lectin. Journal of Clinical Investigation. 2020;130(9):4811–4830. doi:10.1172/JCI134214.

41. Young SAE, Rummler M, Täieb HM, Garske DS, Ellinghaus A, Duda GN, et al. In vivo microCT-based time-lapse morphometry reveals anatomical site-specific differences in bone (re)modeling serving as baseline parameters to detect early pathological events. Bone. 2022;161(May):116432. doi:10.1016/j.bone.2022.116432.

42. Young SAE, Heller AD, Garske DS, Rummler M, Qian V, Ellinghaus A, et al. From breast cancer cell homing to the onset of early bone metastasis: the role of bone (re)modeling in early lesion formation. Science Advances. 2024; in press.

43. Lassen NE, Andersen TL, Pløen GG, Søe K, Hauge EM, Harving S, et al. Coupling of Bone Resorption and Formation in Real Time: New Knowledge Gained From Human Haversian BMUs. Journal of Bone and Mineral Research. 2017;32(7):1395–1405. doi:10.1002/jbmr.3091.

