## Supplementary figures and images for "Phase diagrams of bone remodeling using a 3D stochastic cellular automaton"

### S1_Fig

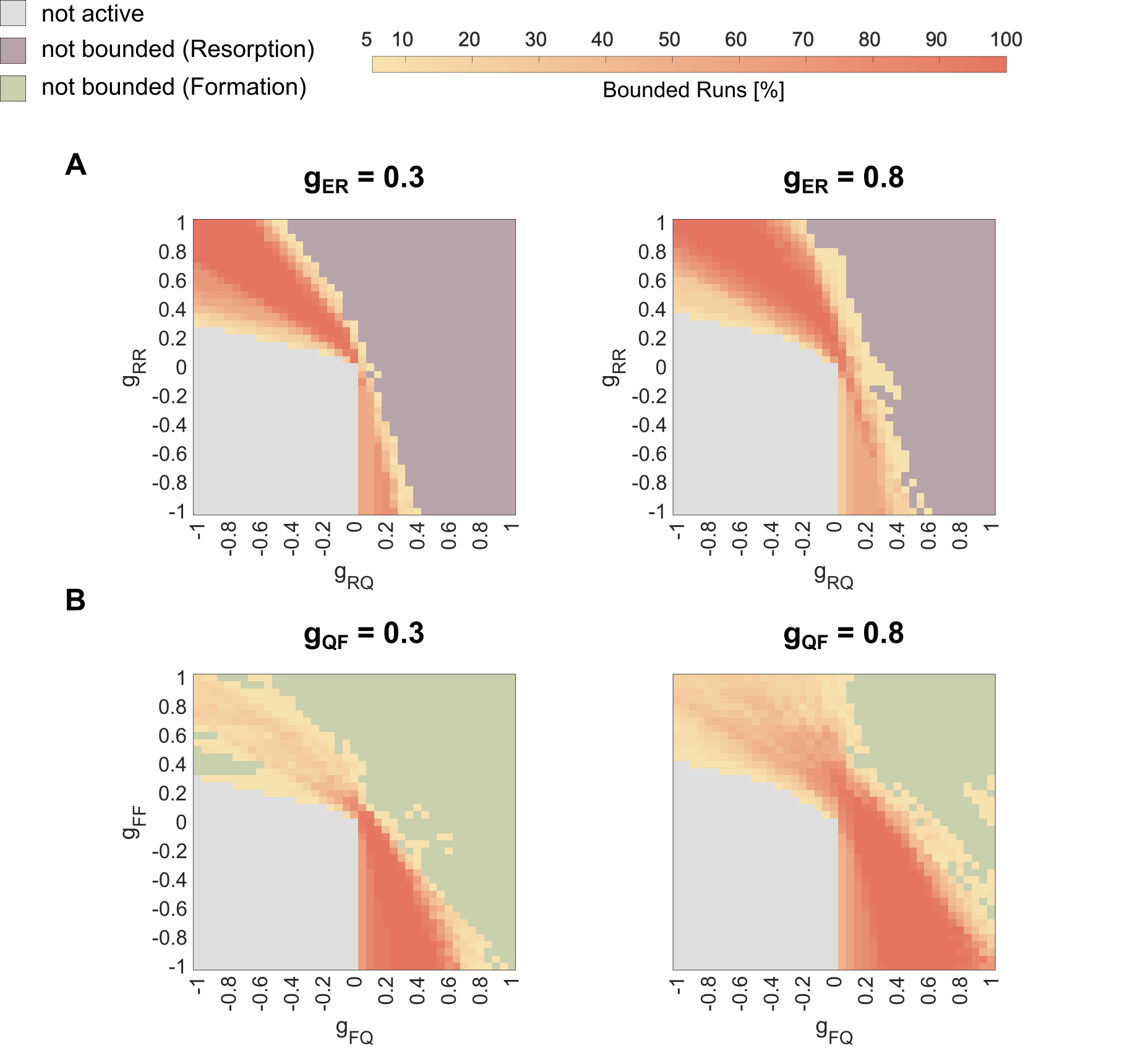

### S2_Fig

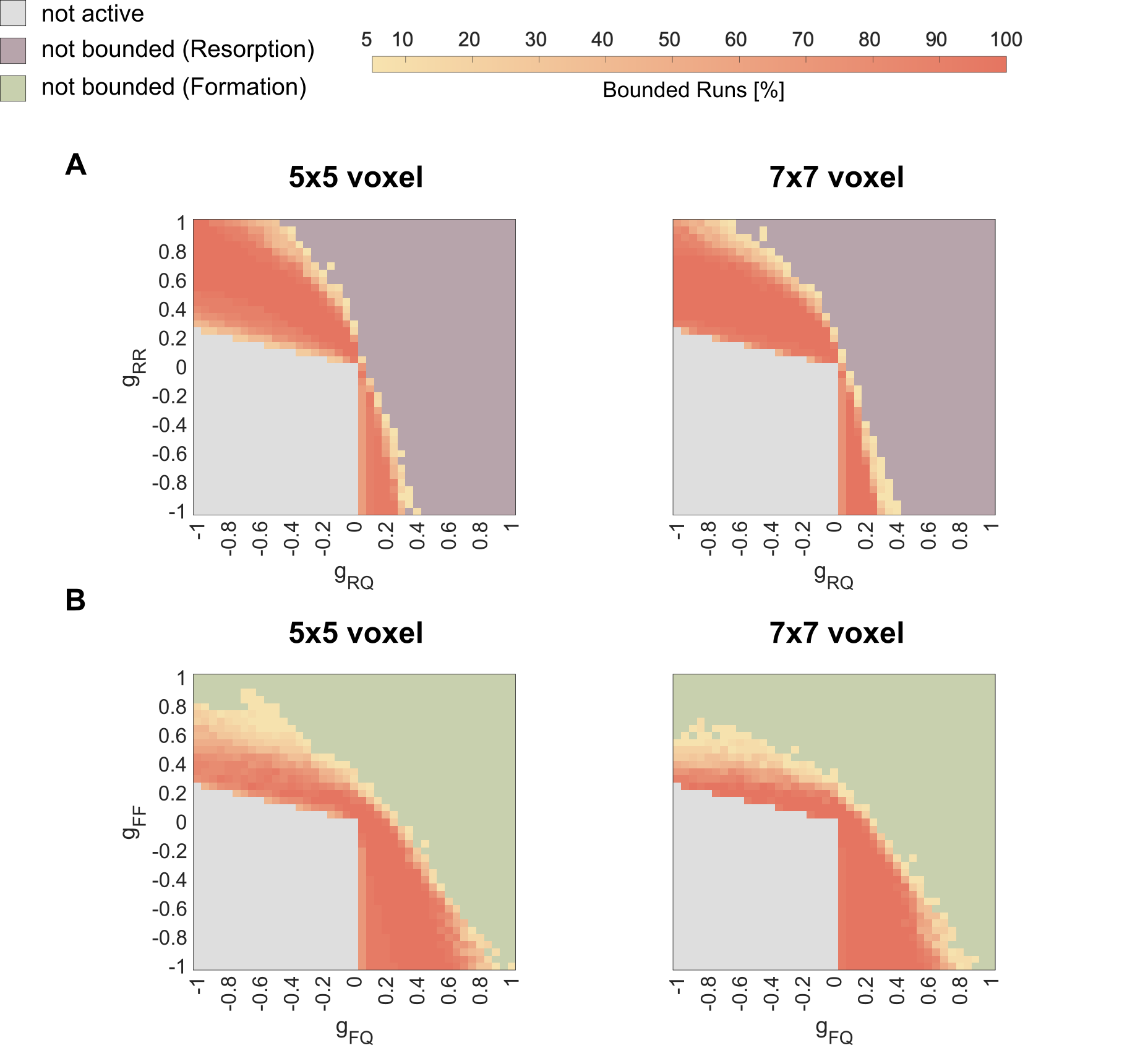

### S3_Fig

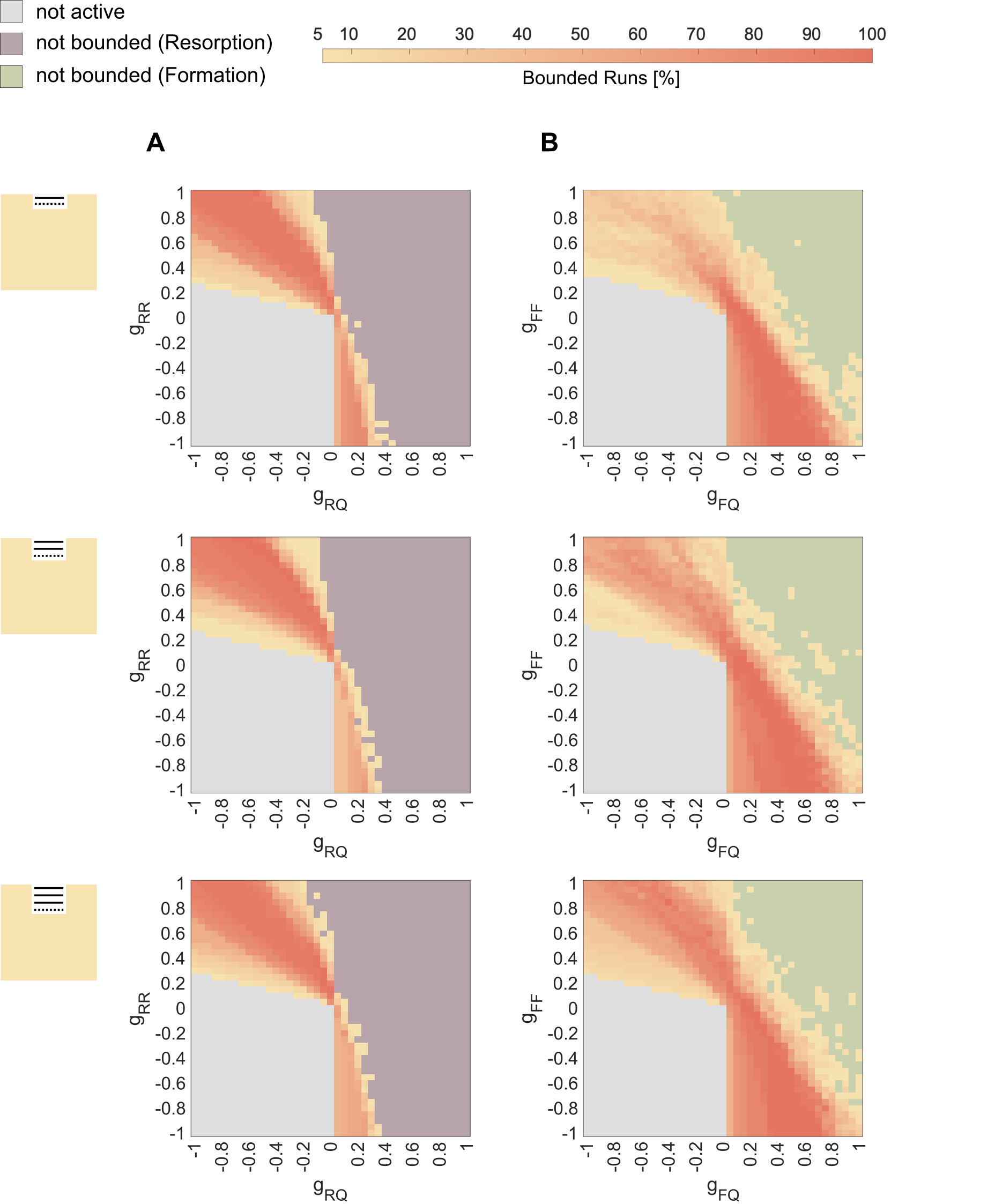

### S4_Fig

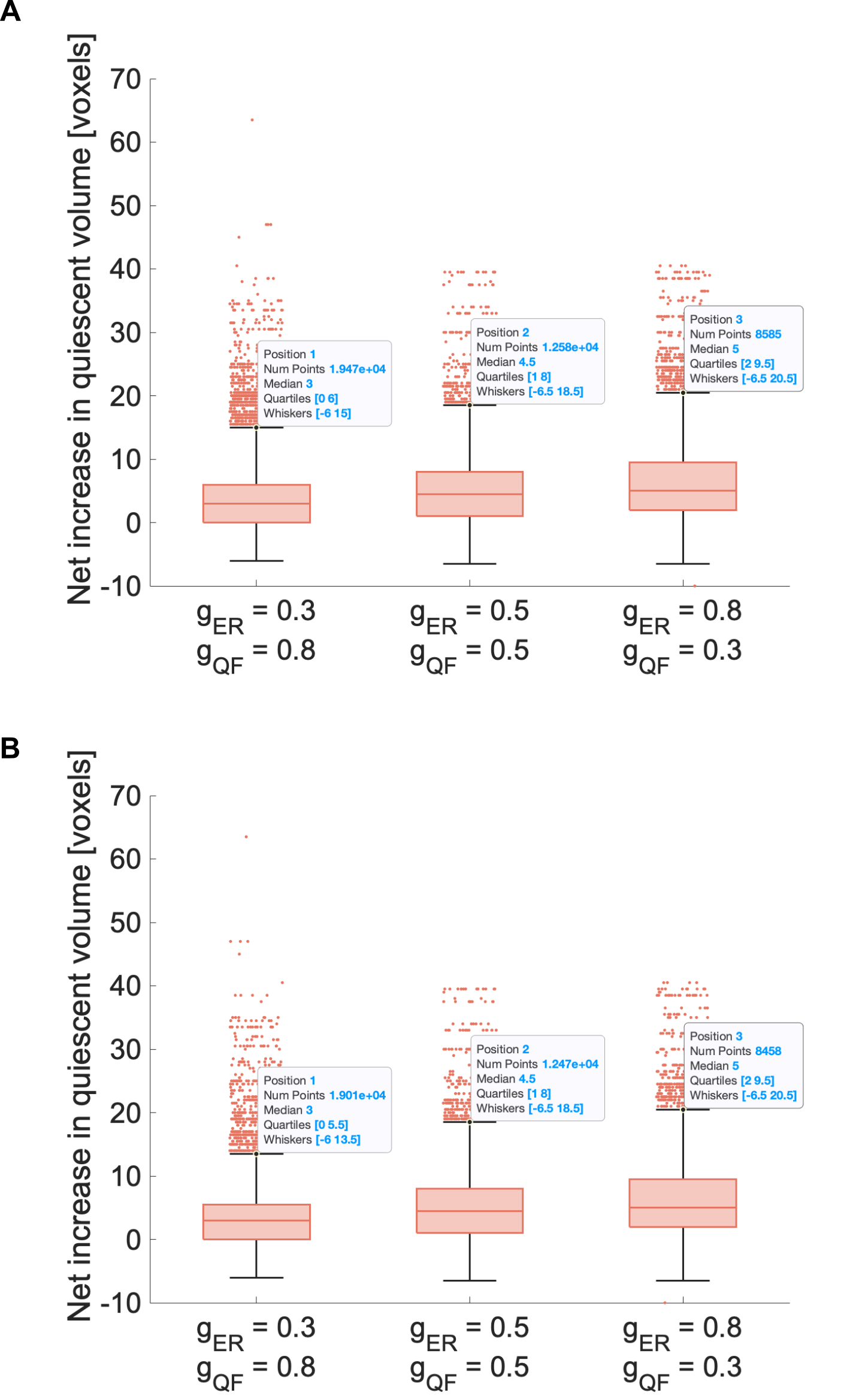
